# An Application for Automated Drosophila Locomotor Assay with Integrated Device Design and Computer Vision Tracking

**DOI:** 10.1101/2025.10.31.685789

**Authors:** Dave Melkani, Neel Harnwal, Shubhankar Desai, Dev Patel, Girish Melkani

## Abstract

*Drosophila* has long served as a powerful model for investigating locomotor behavior, and geotaxis assays have generated valuable insights into genetics, aging, and neurobiology. Nonetheless, their use can be constrained by subjective scoring, modest throughput, and challenges in reproducibility. To complement and extend these classical approaches, we developed and validated an integrated hardware–software platform that enables automated, high-resolution locomotor analysis across 12 vials in parallel. The system integrates 3D-printed mechanical components, Raspberry Pi–based video acquisition, and programmable environmental controls to ensure standardized conditions. A deep learning pipeline segments vials with near-perfect accuracy (IoU > 0.95), while computer vision algorithms quantify climbing trajectories, velocity, and positional zone occupancy at 60 frames per second. The end-to-end workflow converts raw video into time-resolved metrics, supports sex-specific aggregation, and incorporates advanced statistical analyses, including Linear Mixed Effects regression, harmonic mean p-values, and Mann–Whitney U tests. Relative to manual scoring, this automated pipeline yields 2.8-fold faster processing and nearly 800-fold higher data density. Application of the platform uncovered reproducible phenotypes of multiple genotypes. For example, a circadian mutant known as *Clock*^*out*^, males displayed progressive climbing deficits with age, whereas females-maintained age-resilient trajectories. Moreover, male *Clock*^*out*^ exhibited a reduced performance compared to age-matched control (*w*^*1118*^), however, female *Clock*^*out*^ showed subtle reduction in performance. Additionally, glial-specific knockdown of *PolG*, encoding the DNA polymerase gamma catalytic subunit, revealed striking sex-dimorphic aging patterns: females outperformed controls at older age, while males exhibited marked decline. To promote broad adoption, a user-friendly Python interface (Tkinter GUI) enables accessibility independent of computational expertise. Collectively, this standardized, high-throughput framework advances the resolution of genotype-, age-, and sex-dependent locomotor dynamics, offering new opportunities in aging, circadian biology, and neurodegeneration research.

## 1. Introduction

*Drosophila melanogaster*, commonly known as the fruit fly, has long been recognized as a pivotal model organism in the field of behavioral, physiological and cell-molecular genetics. Its utilization in research can be attributed to several key characteristics that make it particularly suitable for studying complex behaviors, including locomotion [1]. Research on geotaxis, which is the orientation of flies’ response to gravity, has been particularly informative in understanding the neural and genetic underpinnings of motor function and behavior. Previous studies have demonstrated that *Drosophila* exhibits robust geotactic responses, which can be quantitatively assessed through a variety of experimental paradigms. Such assessments have revealed insights into the neuromuscular mechanisms and signaling pathways that govern locomotor activities, as well as the effects of genetic mutations and pharmacological treatments on these responses [2].

Traditional geotaxis assays have relied heavily on manual observation, where researchers typically measure the climbing behavior of Drosophila in response to gravity by tapping flies in vials, manually tracking their climbing over a fixed interval, and recording how many reaches a threshold within a set time [1]. While these methods have provided foundational insights into locomotor function, they are inherently limited by subjective interpretation and low throughput. Manual scoring is prone to human error, and the need for repeated observations introduces variability, reducing the reliability of the results. Additionally, the traditional methods often require significant amount of time and labor for counting the flies. The advent of computational ethology provided a paradigm shift in behavioral tracking. The landmark work by Branson et al. introduced a high-throughput, camera-based framework capable of automatically quantifying individual and social behaviors in large groups of Drosophila, establishing the foundation for modern machine-vision approaches to behavioral analysis [3]. As *Drosophila* behavior can be subtle and nuanced, particularly in the context of aging or neurodegenerative studies, the need for more precise and scalable methods has become evident. Automated assays present an appealing solution, offering the potential to capture fine-grained data across multiple experimental replicates and time points with significantly reduced bias. In response, several automated or semi-automated methods have been developed. The RING assay provided a high-throughput but snapshot-based approach to geotaxis [4]. Subsequent iterations integrated automated video capture and analysis pipelines [5,6], while more recent open-source platforms such as FreeClimber [7] and high-resolution video frameworks [8] enable continuous, automated quantification of climbing trajectories. These methods highlight the field’s recognition of the need to replace manual approaches with scalable, less biased tools. However, these tools do not provide the technical and analytical depth that pushes the field towards individualized, statistically rigorous locomotor profiling. These tools also need to enhance the ability to conduct high-throughput experiments. Here, we ask whether a fully automated, vision-based geotaxis assay can (i) reproduce the sensitivity of manual scoring to subtle locomotor phenotypes, (ii) provide a more comprehensive analytical scope than other automated systems, and (iii) enable high-throughput screening of genetic or pharmacological agitations without loss of accuracy.

The geotaxis assay platform combines a custom-built hardware setup featuring high-resolution cameras, lighting arrays, and tight control for environmental variables, with an end-to-end PyTorch (v2.3.1+cu121) deep-learning pipeline. All model code and data handling execute seamlessly on NVIDIA GPUs with CUDA 12.1. We employed the torchvision Faster R-CNN implementation with a ResNet-50 backbone and Feature Pyramid Network, replacing its standard box_predictor head to detect vial versus background. By having the models without any pre-trained weights (it starts with random weights instead of ones it’s trained on), the system is built on top of a custom-trained deep learning model, rather than using an off-the-shelf model trained on unrelated data. This means that the Faster R-CNN model is trained from scratch on the custom dataset of *Drosophila* vial images, allowing the model to learn vial-specific features that are relevant for geotaxis assay setup. This software-hardware pipeline is fully tailored and self-sufficient, not dependent on general-purpose pretrained detectors. In the model, input frames are converted to tensors and processed with the COCO format annotations, enabling on-device batching (batch size 1) and unified GPU acceleration for both training and inference. This tight integration of device and deep-learning software not only automates video capture and data collection but also standardizes boundary detection across trials, minimizing human error and ensuring consistent comparative analyses of locomotor behavior under varying genetic and environmental conditions.

In the current manuscript, we (1) designed and validated an automated geotaxis platform integrating high-resolution imaging and environmental control. (2) Developed and evaluated deep-learning-based and computer-vision-based algorithms for vial segmentation and individual-fly tracking. (3) Benchmark the automated measurements against manual scoring across multiple genotypes and treatment conditions. (4) Demonstrated the system’s utility in a pilot screen for locomotor impairments in aging and mutant models. These aims set the stage for a comprehensive evaluation of our system’s accuracy, robustness, and throughput compared to traditional methods.

## 2. Materials and Methods

### *Drosophila* strains and rearing conditions

As recently reported, *Drosophila* stocks were maintained on a standard cornmeal-yeast-agar diet containing nipagin and propionic acid to prevent microbial growth (ref. All the flies were reared under controlled environmental conditions at 22 °C with 50% relative humidity and a 12:12 hour light-dark circadian cycle, with routine food changes [9]. The following stocks were used. Wildtype stock (w^1118^), Circadian mutant (*Clock*^*out*^) from Bloomington Drosophila Stock Center (BDSC # 24515). To evaluate the locomotor performance of the gene linked with repair of the mitochondrial genome, called DNA polymerase gamma subunit 1 (PolG1) encodes the catalytic subunit of mitochondrial DNA polymerase gamma, RNAi stocks were obtained from Vienna Drosophila Resources Center (VDRC # V106955) and BDSC #31081. Glial-specific knockdown was carried out using Glaz-Gal4 (BDSC# 8765), using the UAS-Gal4 approach as previously reported [10]. Briefly, the progeny of the above-mentioned flies will be collected at eclosion, separated by sex, and held at a density of 25 flies/vial for experimentation as routinely used in the lab.

### Geotaxis Assay Device Development

The geotaxis assay device was built using custom-designed 3D printed parts and an integrated motor system. The motor system, based on a third-class lever, facilitates precise tapping of the vial holders. The recording of the video is done using a Raspberry Pi 4B device and a Raspberry Pi High Quality Camera with a 12.3-megapixel Sony IMX477 sensor. Researchers can use a standard monitor and keyboard to run a python script enabling the camera to record at 60 frames per second at 1280 by 720-pixel resolution and the motor to activate causing the device to tap 3 times (grounding the flies), it then pauses for 15 seconds as the flies try to climb up. This process happens 4 times and then the camera stops recording. Afterwards, the full video is saved as a h264 video file, which researchers can then export using a flash drive as they reduce the size of the video file while maintaining the quality. To ensure video clarity, a light source is directly pointed at the vials, which prevents glare from the above ceiling lights reflecting off the glass vials. This also helps illuminate the flies climbing in front of the white background for more accurate tracking. **Supplementary Figure 1a** shows the full flowchart of this process. The printed vial container can hold a total of 12 vials for researchers to use for different genotype studies. A STL screen capture shown in**Supplementary Figure 1b** displays the vial holder well.

## 3. Results

### Experimental Protocol for Geotaxis Assay

All the video files were saved by h264, and they have provided a metadata comma-separated values (CSV) file for the specific video recorded. This metadata file contains 4 columns: “Vial_Num”, “Genotype”, “Gender”, “N”. “Vial_Num” represents the specific vials that the flies are in within the vial container, “Genotype” is the name of the genotype, “Gender” is the vials sex, and “N” is the number of flies in a vial. The experimental setup is organized in a hierarchical file folder structure as shown in **Figure 1a**. At the top is the main folder, which serves as the root directory of the experiment. The main folder contains multiple subfolders, where each corresponds to a video unit. The number of subfolders can vary based on the specific experiment requirement. Each subfolder is uniquely named based on the h264 video file contained in it. For example, if a video file is named Set1_W1118_04_17_25.h264, the subfolder containing it will be named Set1_W1118_04_17_25. Each subfolder contains two essential files: the h264 video file and the CSV metadata file, consistently named “geotaxis_metadata.csv” across all the subfolders. Once this main folder is populated with the necessary files and folders, it is placed in the same directory as the main Python script for analysis. This script iteratively accesses the subfolders within the main folder, analyzing the videos and outputting the relevant results.

**Figure 1.**
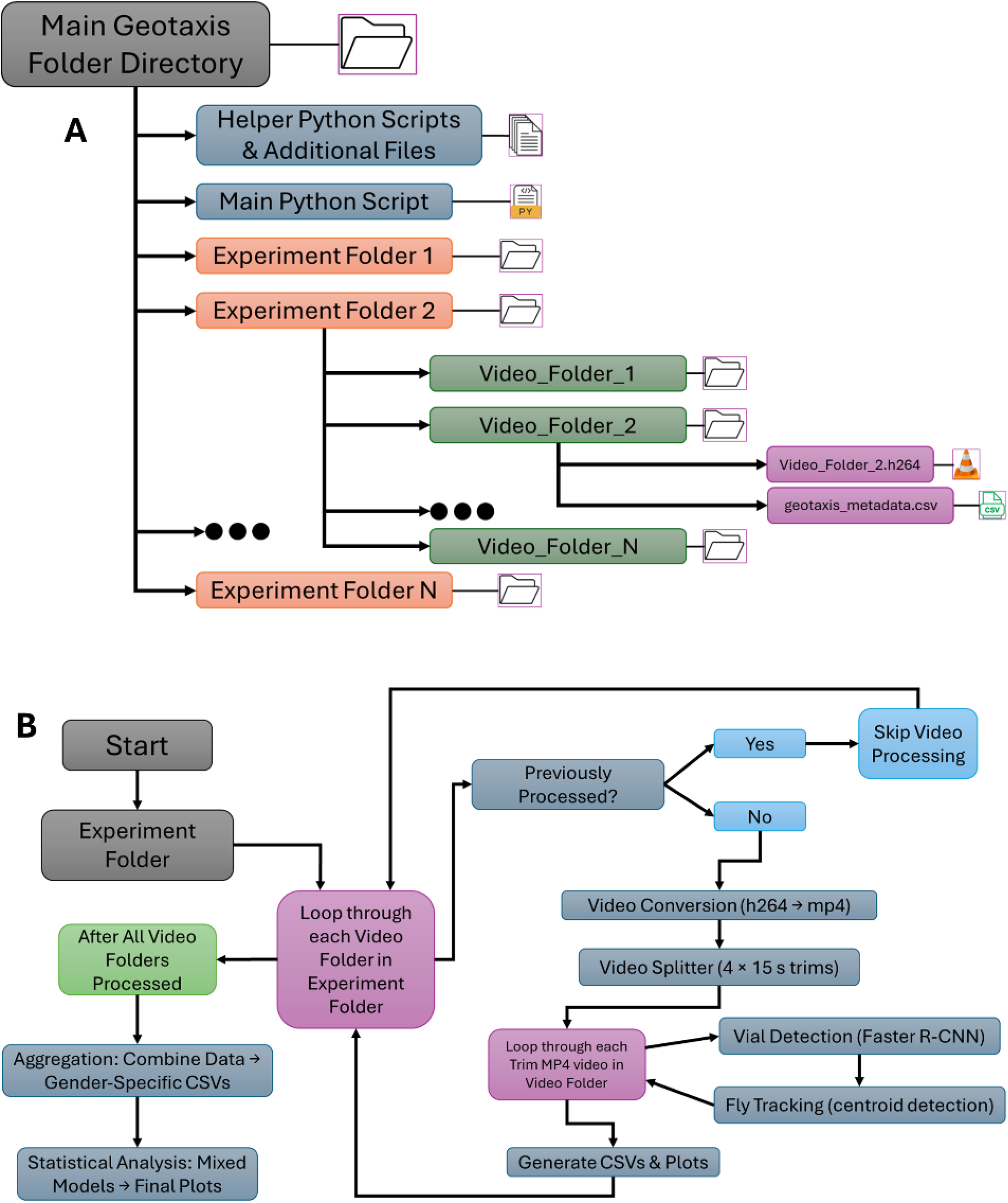
Experimental Workflow and Folder Structure: (a) Directory tree hierarchical diagram of the main script folder, subfolders per video (named by h264 file), and contents (.h264 video + CSV Metadata). (b) Processing workflow chart showing loop over video folders → conversion & splitting → vial detection → fly tracking & plotting → post-iteration aggregation → statistical analysis.

### Behavioral Analysis

This section outlines the methodologies employed for conducting behavioral analysis on the geotaxis data utilizing computer vision techniques and a deep learning model. After acquiring all the relevant data stored in the main folder, researchers initiate the analysis by running a Python application. The app prompts the users to input the main folder that has all the experiment folders where the user will specify their geotaxis experiment that they want to process. The code then parses through each of the subfolders within the input experiment folder and follows a series of steps in the processing. Firstly, it checks if the current subfolder in the iterative process has all the required files, implying that this folder has been previously analyzed. If that subfolder contains all the required files, it skips that folder and moves onto the next. If the next folder does not contain all the required files, it starts the processing pipeline for that video, which are video conversion, video splitter, vial detection, fly detection, plot generator. Finally, after all the video folders within the main folder have been processed, the programs run the experiment aggregator and statistical analysis pipeline across all the video folders. **Figure 1b** illustrates this pipeline loop well.

### Video Conversion

To start, the pipeline first converts the h264 video file into an mp4 video file using an ffmpeg command. It has “qscale=0” for high quality video encoding with minimal compression and “r=0” for ensuring the video outputs at 60 frames per second as default parameters.

### Video Splitter

After the mp4 video is created, this process takes the mp4 video and splits it into four mp4 video each with 15 seconds of recording. With the input mp4 video, it is first processed by detection motion using using “MOG2” which is a function in OpenCV (a Python library used to examine image & video data) that creates a background using the mixture of gaussians algorithm. It then creates a binary mask highlighting moving objects and finds contours, detecting the objects that are in motion. Finally, the contour areas are summed representing total movement while also filtering out small areas less than 2000 pixels. The total movement is saved in a list for each frame of the input video. Using the scipy signals function “find_peaks”, the code looks for significant spikes in the stored list identifying moments of high motion (heights greater than 100,000). Inactivity is defined as motion less than 30,000 and if 270+ consecutive frames are defined as inactive, an end frame is defined but capped to 900 frames (15 seconds). The start and end frame indices for each period of inactivity are saved. With each start and end indices pair, the main mp4 video is trimmed and saved resulting in 4 trimmed mp4 videos. After the video conversion and splitting process, **Figure 2a** shows a detailed flow of each trim video showing the vial and fly detection process for each trim video in each video folder.

**Figure 2.**
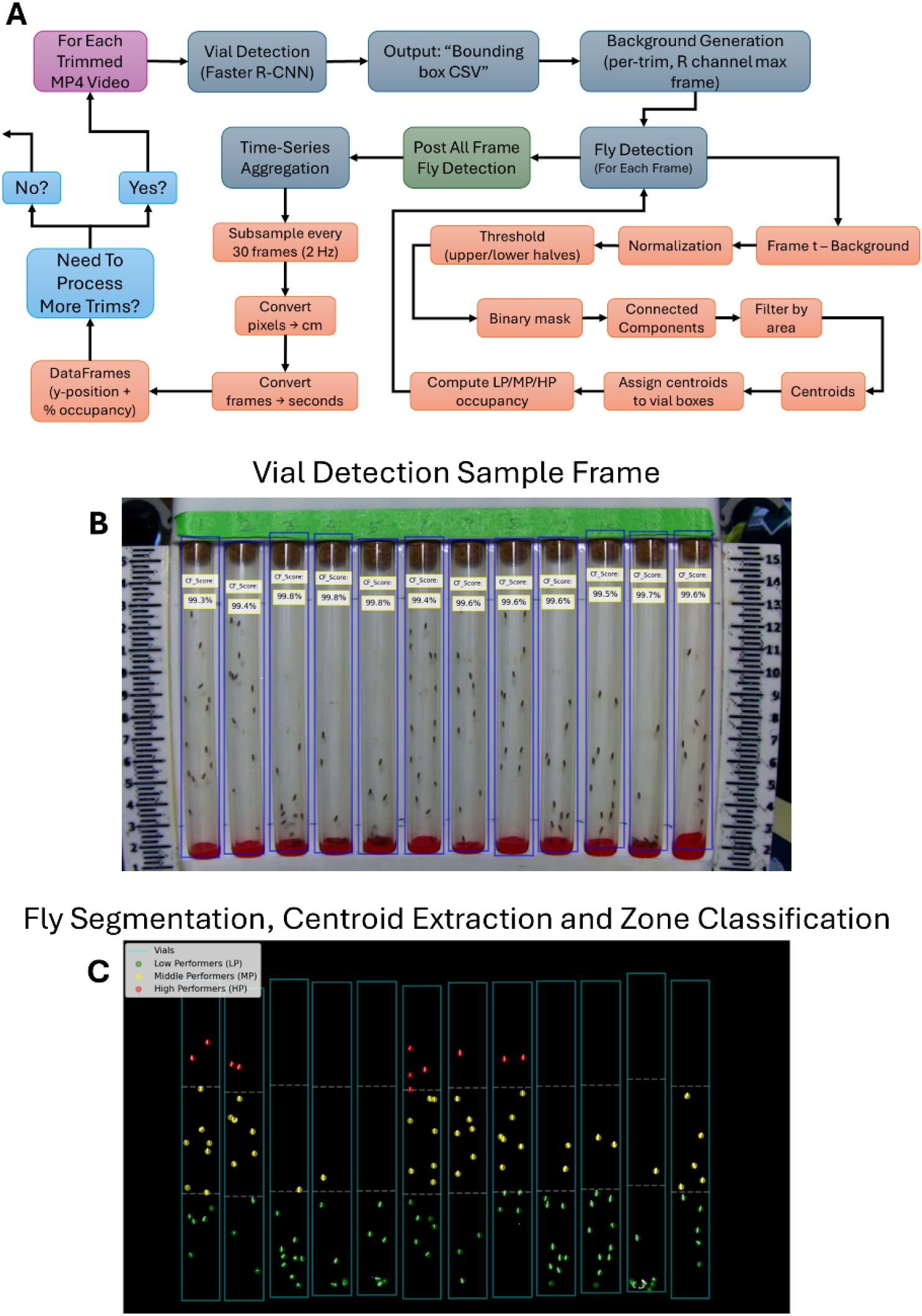
Computer Vision Pipeline for Vial and Fly Detection. (a) High-level box-and-arrow diagram pipeline schematic summarizing the computational workflow for each trimmed video. (b) Sample frame overlaid with Faster R-CNN bounding boxes and labels for each vial, binary mask of moving objects (after background subtraction) with centroids marked for individual flies divided into low (LP), middle (MP), and high (HP) performer zones.

### Vial Detection

For each trimmed MP4 video in a video folder, a single frame is randomly selected, converted from BGR to RGB, wrapped as a PIL Image, and transformed into a tensor for model ingestion. Vial detection leverages a custom VialDataset class that reads 1,140 annotated frames (1280 × 720 pixels) from a COCO JSON file containing 13,680 vial instances (12 vials per frame), assembling bounding boxes, labels, areas, and iscrowd flags for seamless compatibility with torchvision’s detection API. Training runs for one epoch (1,140 iterations) with an SGD optimizer (lr = 0.005, momentum = 0.9, weight_decay = 0.005), achieving rapid convergence (loss falling from ∼2.04 to ∼0.013). Inference filters predictions at confidence ≥ 0.85, sorts the detections left-to-right by x-coordinate, and drops any vials that are not present in the metadata before saving the final positions to CSV. This automated vial-detection step, illustrated in **Figure 2b**, ensures accurate region-of-interest (ROI) selection for subsequent fly-tracking analyses without manual intervention.

### Fly Detection

In this pipeline, once the list of N experiment-specific video folders is generated, each folder is processed in turn by instantiating a custom class object, with parameters for frames per second, a subsampling step size, thresholding values for the upper and lower halves of the chamber, and pixel-based adjustments, pointing to that folder’s path. The run method then first locates the four “trim” MP4 videos, considered each videos technical replicates, and their corresponding vial-position CSVs for the specified video folder; it reads each video fully to compute a background image by taking the per-pixel maximum over all extracted frames, which isolates and normalizes the red color channel. After loading the genotype metadata (which defines vial numbers, expected numbers of flies, and their x/y bounding-box coordinates), the code enters a nested loop over each trim (1–4) and then over the first 780 frames of each trim video. For each frame, the live-video image (again using the red channel) is subtracted against its trim-specific background, normalized, then threshed separately in the top and bottom halves to segment moving flies into a binary mask. Connected components in this mask are identified and filtered by area to yield centroid points corresponding to individual flies. Each centroid is then spatially assigned to the correct vial by checking which bounding box (from the vial CSV) contains it. Within each vial, centroids are further classified into three vertical zones: high performer (HP), middle performer (MP), & low performer (LP), based on the vial’s y-coordinate boundaries, with special handling of early frames and a y1-based adder threshold to ignore false positives that delay counting MP & HP until the flies are fully in view. Next, **Figure 2c** shows the individually tracked flies of a single frame along with which zone the flies are in for each vial. For each zone and vial at each frame, the code records raw x/y lists and counts, computes percent occupancy of each zone, and stores these in nested dictionaries. Once all frames are processed, the script subsamples by every 30 frames (2 Hz) to build Pandas (a Python data organization library) DataFrames of converted y-positions and percent occupancy for each zone. To convert pixel distances to physical distances, each adjusted y-displacement is scaled according to the chamber holder height (17 cm) relative to the image resolution (720 pixels), providing centimeters per vial position 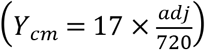. Similarly, frame indices are divided by the camera’s frames-per-second (fps) rate to obtain real-time values in seconds 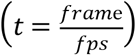, ensuring both spatial and temporal measurements are biologically interpretable. These per-trim DataFrames are then aggregated across the four trims by calculating means and standard errors over replicates for each frame and vial (ordered according to the genotype metadata), yielding summary tables of geotactic performance over time. Finally, the method writes out CSVs and produces quality line plots with error bars showing ±SEM across 13 seconds to visualize how each genotype’s flies climb and redistribute among low, medium, and high zones over the course of the trial, thereby quantitatively capturing the biological geotaxis behavior under study.

### Experiment Aggregator

In this follow up pipeline, all aggregated per-trim CSV outputs generated by the custom class processing are loaded from each experiment subfolder, namely the genotype metadata and the four zone-specific summary files (the total position csv for overall climbing height and the LP, MP, and HP percentage csv files for low, middle, and high zone occupancy), into five parallel lists of DataFrames. The script then creates separate output directories for male and female cohorts, reflecting the necessity of sex-specific behavioral analysis. The core function zips these lists and iterates through each set of metadata and measurement DataFrames: for each vial in the genotype sheet, it assigns a unique replicate label (e.g., “GenotypeA_rep1”) by counting how many times that genotype has appeared for a given sex, then extracts the time-series of mean climbing height and mean percent occupancy for LP, MP, and HP zones by matching vial identifiers. Empty subsets trigger warning prints to flag missing data. These series are stored in nested dictionaries keyed first by sex (“M” or “F”), then by replicate label. After processing all subfolders, these dictionaries are converted into wide DataFrames where rows are replicates and columns are time points (seconds), sorted numerically. Next, a velocity DataFrame for each sex is computed by taking the numerical gradient of climbing height against time, capturing the instantaneous climbing speed (cm/sec). All ten resulting DataFrames (male/female × position, velocity, LP, MP, HP) are then written as CSVs into their respective sex-specific output folders. Finally, a plotting method reshapes each DataFrame into an aggregated time-axis index, groups the replicates by genotype, and computes for each time point the mean, standard error of the mean (SEM), and replicate count (N), thereby yielding aggregated genotype performance curves. Error-bar plots are produced with embedding N in the legend, and tailored axis labels so that male and female geotaxis behaviors (climbing height, speed, and zone occupancy) can be directly compared across genotypes over time. This is shown clearly in **Figure 3** for all the 5 outputs: position (cm), velocity (cm/sec), LP (%), MP (%), and HP (%).

**Figure 3.**
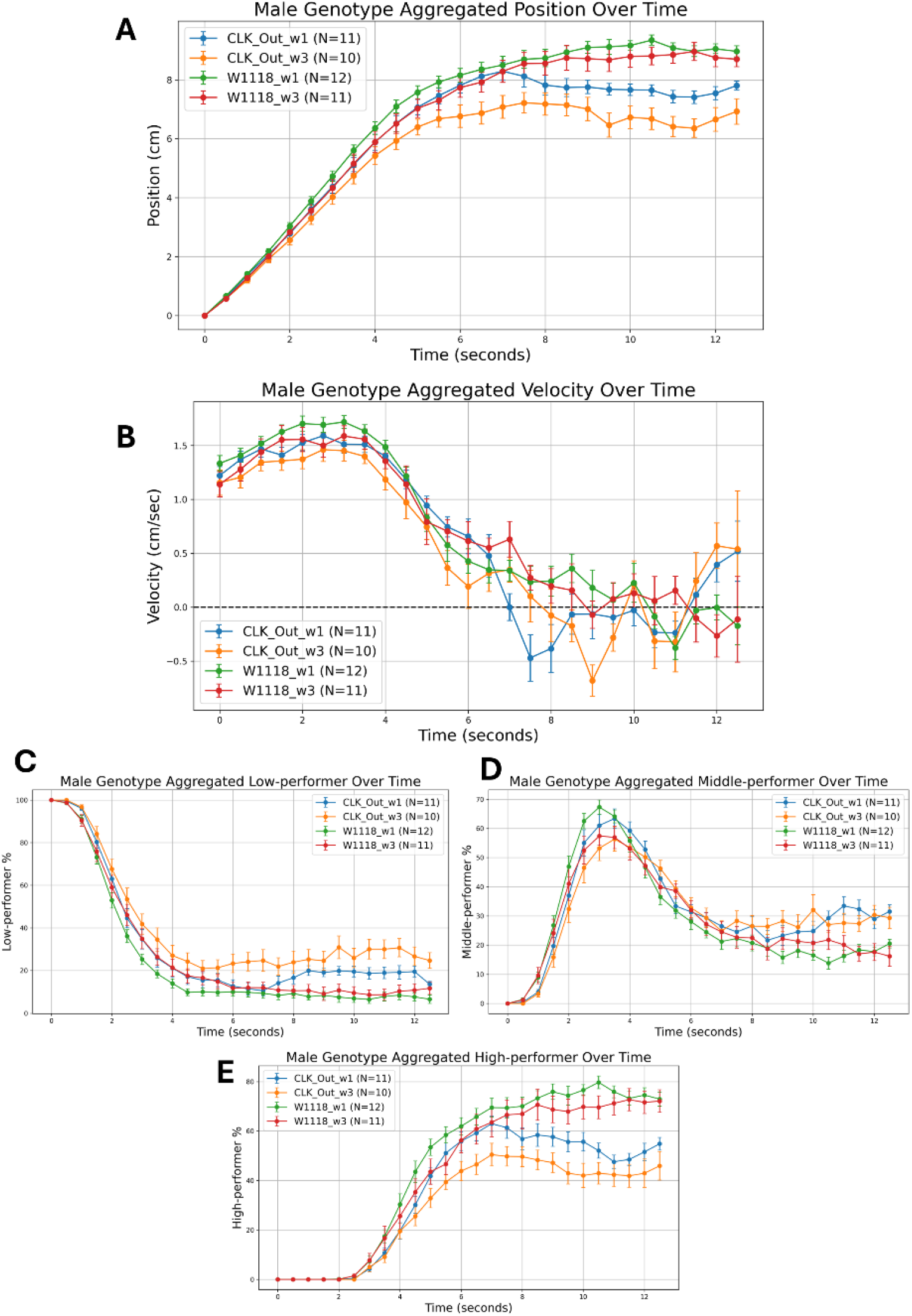
Representative Climbing Trajectories and Zone Occupancy. (a) Mean climbing height (cm) ± SEM and (b) mean climbing velocity (cm/sec) ± SEM over a 13-second recording. These trajectories illustrate consistent upward movement and varying speed dynamics captured across all replicates. (c) Mean low performers (LP) percentages ± SEM, (d) Mean middle performers (MP) percentages ± SEM, and (e) Mean high performers (HP) percentages ± SEM over a 13-second recording. This panel demonstrates temporal changes in population-level zone occupancy based on vertical position thresholds.

### Statistical Analysis

After aggregating across all biological replicates, the DataFrame for each sex (male and female) and each data modality: position (cm), velocity (cm/sec), low performer (%), middle performer (%), high performer (%), is cleaned into a long-form time-series DataFrame denoted as *D* ⊂ ℝ^*n* × 5^ where each row is an instance of the columns: Genotype *G*_*i*_ ∈ {*G*_0_,*G*_1_,…,*G*_*k*_} (categorical variable), Replicate ID *R*_*j*_ (subject identifier), Time *t*, Measurement *y* ∈ ℝ (either position, velocity, or performance metric), and Subject ID *S*_*ij*_ = “*G*_0_” + “_” + “*R*_*j*_”. The data is then filtered to only include the control genotype *G*_*c*_, the selected comparison genotypes *G*_*f*_, …, *G*_*l*_ (where *c* ∈ *G*, 0 ≤ *f and l* ≤ *k*), and only observations where time *t* ≤ *T*_max_.

A Linear Mixed Effects (LME) model is then applied to the long-form data for each modality. LME models provide a statistically principled framework for analyzing longitudinal or clustered data where repeated measurements introduce within-subject correlations. Unlike standard linear regression, LMEs allow the simultaneous modeling of fixed effects, which represent population-level influences, and random effects, which capture subject-specific deviations or trajectories. This structure enables valid inference in the presence of errors that can be correlated, unbalanced designs, and hierarchical data which are conditions that are typically present in behavioral and biological assays. The theory behind mixed models has been extensively researched and developed, offering rigorous approaches to handle repeated measures and longitudinal dependencies [12, 13]. West et al. [14] and Verbeke et al. [13] emphasize their flexibility for modeling both continuous and discrete longitudinal outcomes, while Edwards et al. [11] proposed an *R*^2^ statistic for assessing the explanatory power of fixed effects in such models. Gibbons et al. [12] discuss extensions of these frameworks to multi-level and multivariate data, showing their utility in behavioral research where outcomes are temporally structured. The present analysis adopts these established principles within a modern computational environment using Statsmodels, a Python library that provides a high-level interface for fitting LME models via Restricted Maximum Likelihood Estimation (REML), ensuring unbiased estimation of variance components [11, 14]. The purpose of the LME model in this paper is to model and test for the fixed effects of genotype and time, along with random effects due to repeated measurements from the same subjects (replicates), up to a user-defined time cutoff *T*_max_. The model is constructed as:

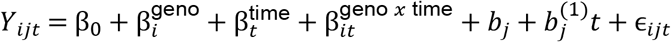

Where *Y*_*ijt*_ measurement for genotype *G*_*i*_ replicate *R*_*j*_ at time *t, β*_0_ is the fixed intercept, 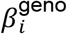 is the fixed effect of genotype *G*_*i*_ (with *G*_0_ as baseline, so for *i* = 0, this is 0), 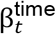 is the fixed effect of time *t*, treated as categorical, 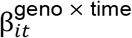 is the fixed effect interaction between genotype and time, 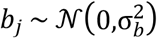 is the random intercept for subject 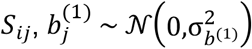 is the random slope for time within subject, and *ϵ* is the residual error. This model was implemented in Python using the *Statsmodels* library to create a formula interface as:

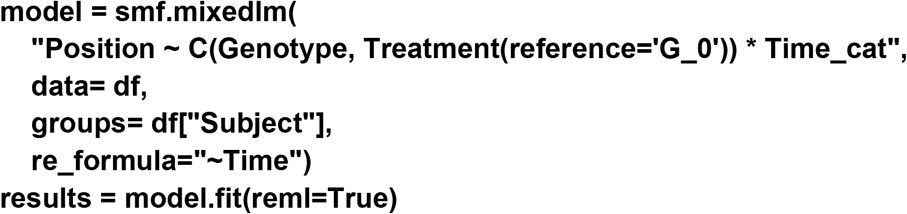

where the random structure *groups=df[“Subject”]* models the correlation between repeated measurements on the same fly (replicate), and *re_formula= “∼Time”* allows each subject to have a random slope and intercept concerning time. The model is fit using Restricted Maximum Likelihood Estimation (REML), which is a standard approach for LME models proving unbiased estimates of variance components, the design matrix for fixed effects, and the design matrix for random effects. After fitting the model, the p-values are extracted for the main genotype effects and genotype-time interactions for all time *t*. These are evaluated using standard Wald tests, and results are annotated with stars based on significance level: Significance = **\*\*\*** for *p* ≤ 0.001, **\*\*** for *p* ≤ 0.01, **\*** for *p* ≤ 0.05, **NS** for *p* > 0.05. Since multiple p-values are associated with each comparison genotype *G*_*m*_ (main effect + all interaction terms), the Harmonic Mean p-value (HMP) method [15] is used to combine them. With *p*_1_,…,*p*_*r*_ being the p values for genotype *G*_*m*_, then:

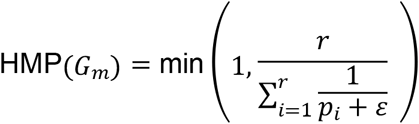

where *ε* = a tiny positive constant to avoid division by zero and *r* is the number of terms (one main effect + multiple interaction terms). Note that this correction method is conservative, controlling family-wise error while combining dependent p-values. This pipeline enables robust detection of temporal genotype-specific effects in high-resolution behavioral time-series while properly modeling within-subject dependencies. These results are added to plot legends that embed replicate counts and p-values, allowing immediate interpretation of statistical robustness alongside sample size. Finally, a line plot is generated showing the mean climbing height over time with shaded error bands (SEM) for each genotype, reference lines at time zero and the cutoff, and axis labels. This final figure is then saved as a PNG file for statistical analysis visualization. Additionally, the Mann-Whitney U Test, a non-parametric alternative to the independent samples t-test utilized to compare median ranks among two genotypes, was employed to complement the mixed-effects linear model by providing independent validation of genotype-level differences at all time points, separate from time-series trends. Its application is justified by its lack of assumptions regarding normality or homoscedasticity.

### Development of Python-Based Application

In addition, we developed a Python application to make the analysis of recorded trials more accessible in a laboratory setting. The tool is designed to eliminate the need for terminal interaction and Python environment configuration so that this method can be employed by those with limited software experience. Interfacing with the application allows the user to process H264 (and MP4) videos, run statistical tests on specified parameters, and produce visualizations of fly position, velocity, and percentage LP/MP/HP via the straightforward workflow described in **Figure 1b**. This application utilizes the Tkinter graphical user interface (GUI) library to enable widget and file dialog functionality. With PyInstaller, a command-line utility for freezing Python programs, we made executables for multiple operating systems that can be launched as an executable from within a folder with the code’s dependencies. The source code is structured such that each screen is a separate Python file which outsources computationally heavy logic for video conversion, vial prediction, fly detection, and statistical analysis from custom helper classes. OpenCV is used to examine frames from the video and track flies with the connected components method. Additionally, Pandas, a Python data organization library, is used to create comma separated value (CSV) files that are plotted with Matplotlib and Seaborn, two data visualization libraries. The application opens to a menu, where the user can select the directory in which they store their recordings and choose the project to be processed. Once entered, the program will convert and trim the videos automatically to track the flies and calculate dependent variables. A log at the top of the menu updates the display with information on the execution of intermediate steps and current video being handled. Raw data obtained from this preliminary analysis is saved to CSVs and graphs in the “Output Males” and “Output Females” folders for the given project. Upon completion, the user can run a Mann Whitney U-test on an input control group and at least one comparison group to yield significance for the five dependent variables throughout a given time range, automatically separated by sex. Data for each test is automatically collected in the output folders, which can be decompressed for easier portability. This process is illustrated in **Figure 4**.

**Figure 4.**
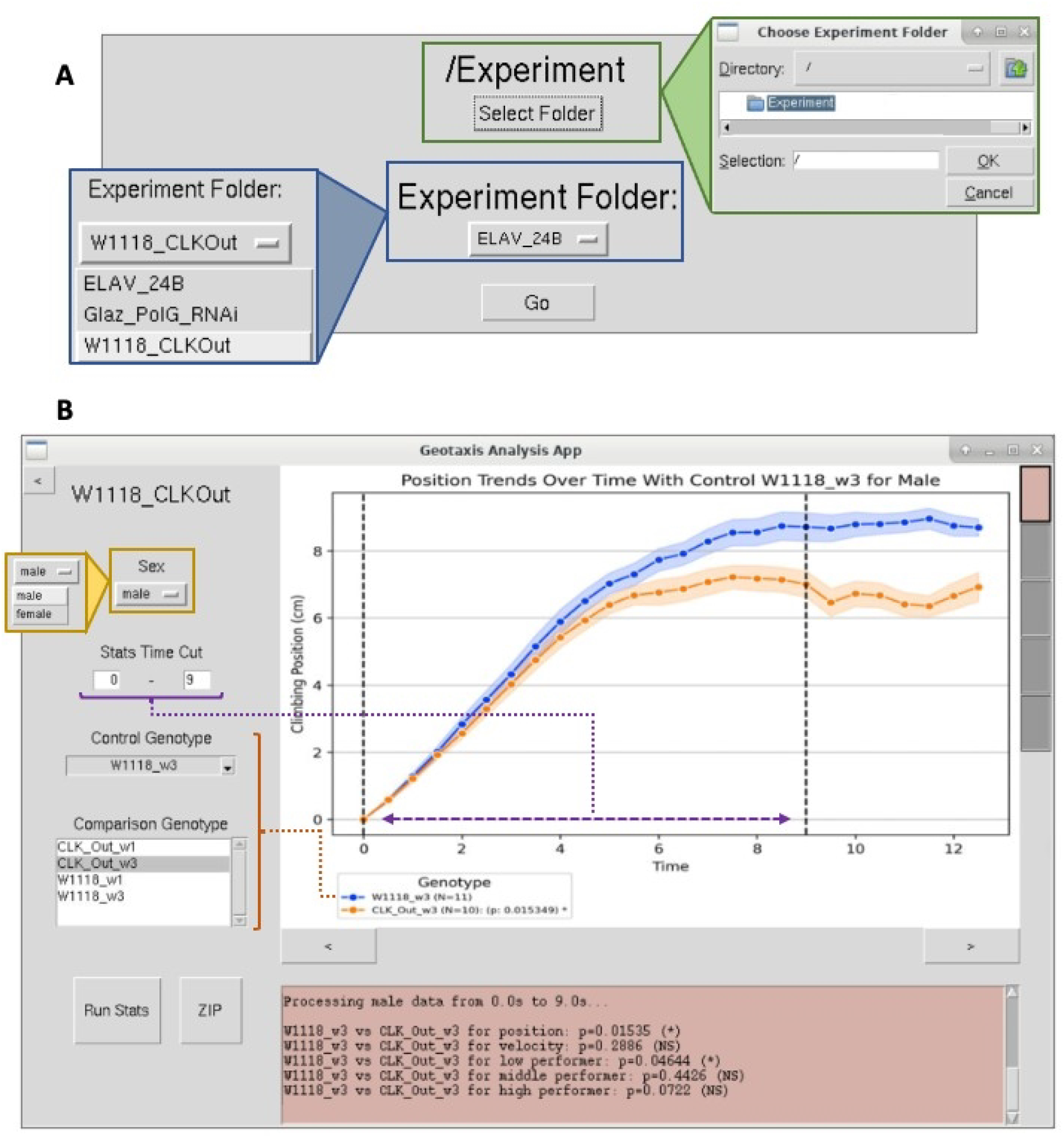
Geotaxis Analysis Linux Application Interface: (a) A view of the menu screen, where the Geotaxis Folder and Experiment Folder are set. (b) A view of the statistical analysis screen, where a display is updated based on specified variables. The analysis screen opens after the user finishes inputting their desired settings on the menu and presses “Go.”

### Clock^Out^mutant showed compromised and progressive geotaxis performance compared to the age-matched control

For assessing strain-specific differences in geotaxis behavior, we applied the automated geotaxis pipeline to probe how disruption of the clock gene affects geotaxis by comparing wild type *w*^1118^) flies with clock out (*Clock*^*Out*^) mutants at two ages: 1 week (wk1) and 3 weeks (wk3). We conducted 4 pairwise comparisons: wk1 *w*^1118^ and wk1 Clock^*Out*^ flies, wk3 *w*^1118^ and wk3 Clock^*Out*^ flies, wk1 and wk3 *w*^1118^ flies, & wk1 and wk3 Clock^*Out*^ flies, for both males and females. **Figure 5** illustrates the results of the 4 comparisons for just male data. For each of the 4 comparisons, this figure shows the mean climbing trajectories (±SEM) with significance from the LME model, a log scaled bar plot of the p values of the remaining 4 LME metrics (velocity, LP, MP, & HP), peak climbing heights of all biological replicates with Mann–Whitney U test annotations, and finally presents a heatmap of time-specific −log_10_(p) from Mann–Whitney U tests p-values, highlighting when each strain diverges from its control. This multi-faceted approach, supported by robust biological replication (N ≥ 10 per genotype), enables detailed dissection of how aging and clock gene disruption affect climbing behavior in Drosophila.

**Figure 5.**
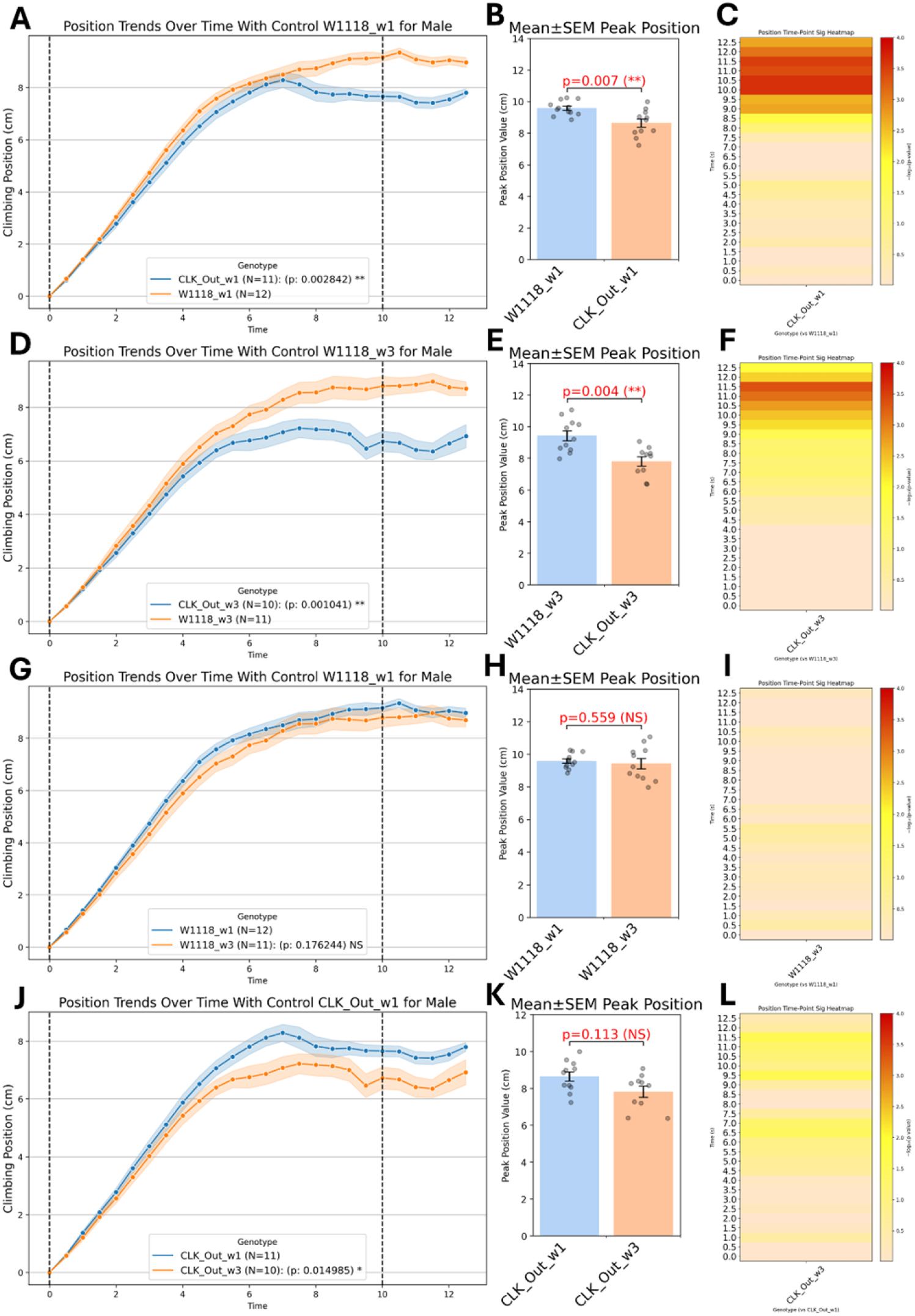
Genotype-Specific Clock Out (Clock^*Out*^ ) Geotaxis Behavior and Statistical Analysis Visualizations. (a) A comprehensive summary of genotype-specific differences in male geotaxis behavior across four pairwise comparisons: genotype effects: w^1118^ wk1 vs Clock^*Out*^ wk1 (A–C), w^1118^ wk3 vs Clock^*Out*^ wk3 (D– F), aging effects: w^1118^ wk1 vs w^1118^ wk3 (G–I), and Clock^*Out*^ wk1 vs Clock^*Out*^ wk3 (J–L). Each comparison includes four subpanels: (1) average climbing trajectories over time (±SEM) annotated with significance from a LME model, (2) a bar plot of individual peak climbing metrics with Mann–Whitney U test annotations; and (3) a heatmap visualizing time-resolved –log_10_(p) values from Mann–Whitney U tests, indicating the temporal windows during which each genotype diverges significantly from its control.

### Mean Climbing Trajectories (LME Model)

#### Clock^*Out*^ males declined with climbing performance and age, while *w*^1118^ stayed stable, and females showed mild aging without genotype effects

For males, the LME model on position trajectories revealed that aging from wk1 to wk3 for the *w*^1118^ produced a non-significant drop in harmonic mean position (p=0.179, NS) which indicates minimal decline across trials for *w*^1118^ aging. However, the Clock^*Out*^ mutants declined significantly in climbing performance from wk1 to wk3 (p=0.0150, *), suggesting a progressive aging effect of clock disruption on climbing over trials. For wk1 flies, we see that the Clock^*Out*^ males climbed significantly lower than *w*^1118^ controls (p=0.0086, **) with a similar pronounced deficit in climbing trajectory persisting for wk1 flies (p=0.0010, **). This indicates a significant genotypic effect in climbing performance at both ages. For females, aging from wk1 to wk3 induced a small but significant decline in harmonic mean position (p=0.0175, *) for *w*^1118^ flies, indicating reduced overall climbing behavior. Aging, however, did not seem to statistically impact the Clock^*Out*^ flies (p=0.303, NS). Additionally, genotype differences between *w*^1118^ and Clock^*Out*^ showed no significant trajectory difference with wk1 flies (p=0.218, NS) and wk3 flies (p=0.770, NS).

### Peak Climbing Height

#### Clock^*Out*^ mutants had lower peak climbing heights than *w*^1118^ in some groups, with no aging effects observed

For males, the Mann-Whitney U test revealed a significant reduction in maximum height for wk1 Clock^*Out*^ flies compared to the *w*^1118^ (p=0.015, *). Peak heights also differed significantly between genotypes for wk3 flies (p= 0.004, **). This shows that even accounting for each replicates peak climbing height, there is a clear statistical difference between the clout out male mutants and the *w*^1118^ male flies for both ages. Peak height, however, did not statistically differ in age comparisons between wk1 and wk3 for *w*^1118^ (p=1.000, NS) and Clock^*Out*^ mutants (p=0.113, NS). For females, wk1 Clock^*Out*^ flies reached statistically lower peaks than wk1 *w*^1118^ flies (p=0.035,*), but the wk3 flies for *w*^1118^ and Clock^*Out*^ comparisons did not differ statistically (p=0.972, NS). Similarly for aging comparisons, the wk1 & wk3 flies *w*^1118^ flies showed no significant aging effect (p=0.135, NS) and the Clock^*Out*^ mutants also showed no difference in peak climbing height (p=0.470, NS).

### Performer Categories (Velocity, LP, MP, HP)

#### Clock^*Out*^ mutants showed more low performers and fewer high performers than *w*^1118^, especially in males, while aging mainly affected *w*^1118^ low performance

For male velocities, all four pairwise comparisons of maximum velocity were non-significant (peak p ≥ 0.275, NS), and the LME harmonic-mean velocity trajectories likewise showed no differences (p = 0.712 for age; p = 0.748 for genotype, both NS). Similarly, no female velocity comparisons reached significance (peak p ≥ 0.164, NS; LME p = 0.215 for age; p = 0.451 for genotype, both NS). In male low performers, the Clock^*Out*^ mutants exhibited a modest but significant increase in LP harmonic means compared to the *w*^1118^ flies for wk1 (LME p = 0.015, *) and wk3 (LME p = 0.020, *). Peak low performers (lowest percentage) at wk3, Clock^*Out*^ mutants tend to have statistically higher percentage of low performers than *w*^1118^ flies (peak p = 0.008, **) with wk1 showing a similar pattern of Clock^*Out*^ mutants having a higher percentage of low performers though not statistically significant (peak p=0.051, NS). Aging effects seem to exhibit in the *w*^1118^ flies from wk1 to wk3 (LME p=0.041,*) but not in Clock^*Out*^ mutants from wk1 to wk3 (LME p=0.078, NS). Peak aging shows no significance between the *w*^1118^ ages and the *Clock*^*out*^ ages. In female low performers, genotype differences showed significance at wk1 (LME p=0.020, *) but not at wk3 (LME p=0.698, NS) with Clock^*Out*^ mutants having higher percentages of low performers. Peak genotype differences showed no significance for either age of the genotype comparisons. Aging effect for *w*^1118^ flies showed significant differences with wk3 *w*^1118^ flies spending more time in low performance compared to wk1 *w*^1118^ flies (LME p=0.0004, ***) while aging effects for Clock^*Out*^ mutants from wk1 to wk3 showed no significance (LME p=0.260, NS). Peak aging differences showed no significance for either genotype of age comparisons. For both male and female middle performers, neither age nor genotype influenced MP. All LME harmonic-mean p values were p ≥ 0.324 (NS) and all peak MP had p values p ≥ 0.117 (NS). For male high performers, Clock^*Out*^ mutants tend to have fewer high performers compared to *w*^1118^ flies at both wk1 (LME p=0.038, *) and wk3 (LME p=0.019, *). Peak high performers also showed a significant difference between the Clock^*Out*^ mutant and *w*^1118^ at both wk1 (peak p=0.001, **) and wk3 (peak p=0.007, **). This shows that male *w*^1118^ flies tend to have significantly more high performers than the Clock^*Out*^ mutants at both ages. Aging effects between the wk1 and wk3 flies seemed to not show significance for both the *w*^1118^ and Clock^*Out*^ (all LME harmonic-mean p ≥ 0.085, NS; all peak p ≥ 0.053, NS). For female high performers, we see no significance in genotype effects at both ages, and we see no significance in aging effects for both genotypes (all LME harmonic-mean p ≥ 0.069, NS; all peak p ≥ 0.098, NS). All the p values referenced can be found for males (**Table 1a**) and females (**Table 1b**).

**Table 1.** **P-values for trajectory and peak metrics comparing w**^**1118**^**/**Clock^***Out***^ **driven lines** for (a) males and (b) females. Footnote: NS, not significant (p > 0.05); * (p ≤ 0.05); ** (p ≤ 0.01); *** (p ≤ 0.001)

| Metrics | W1118_w1 vs CLK_Out_w1 | W1118_w3 vs CLK_Out_w3 | W1118_w1 vs W1118_w3 | CLK_Out_w1 vs CLK_Out_w3 |
| --- | --- | --- | --- | --- |
| Gender | male | male | male | male |
| position_LME | 0.003 (**) | 0.001 (**) | 0.176 (NS) | 0.015 (*) |
| position_Peak | 0.007 (**) | 0.004 (**) | 0.559 (NS) | 0.113 (NS) |
| velocity_LME | 0.209 (NS) | 0.302 (NS) | 0.346 (NS) | 0.235 (NS) |
| velocity_Peak | 0.166 (NS) | 0.275 (NS) | 0.829 (NS) | 0.647 (NS) |
| low performer_LME | 0.015 (*) | 0.023 (*) | 0.041 (*) | 0.079 (NS) |
| low performer_Peak | 0.051 (NS) | 0.008 (**) | 0.579 (NS) | 0.084 (NS) |
| middle performer_LME | 0.267 (NS) | 0.435 (NS) | 0.352 (NS) | 0.417 (NS) |
| middle performer_Peak | 0.310 (NS) | 0.170 (NS) | 0.186 (NS) | 0.275 (NS) |
| high performer_LME | 0.015 (*) | 0.019 (*) | 0.274 (NS) | 0.085 (NS) |
| high performer_Peak | 0.001 (***) | 0.007 (**) | 0.622 (NS) | 0.053 (NS) |

**Table 1. P-values for trajectory and peak metrics comparing $w^{1118}/Clock^{Out}$ driven lines for (a) males and (b) females. Footnote: NS, not significant ( $p > 0.05$ ); \* ( $p \leq 0.05$ ); \*\* ( $p \leq 0.01$ ); \*\*\* ( $p \leq 0.001$ )**
| Metrics | W1118_w1 vs CLK_Out_w1 | W1118_w3 vs CLK_Out_w3 | W1118_w1 vs W1118_w3 | CLK_Out_w1 vs CLK_Out_w3 |
| --- | --- | --- | --- | --- |
| Gender | female | female | female | female |
| position_LME | 0.215 (NS) | 0.770 (NS) | 0.016 (*) | 0.303 (NS) |
| position_Peak | 0.018 (*) | 0.972 (NS) | 0.067 (NS) | 0.470 (NS) |
| velocity_LME | 0.448 (NS) | 0.348 (NS) | 0.208 (NS) | 0.581 (NS) |
| velocity_Peak | 0.082 (NS) | 0.972 (NS) | 0.306 (NS) | 0.896 (NS) |
| low performer_LME | 0.020 (*) | 0.698 (NS) | 0.000 (***) | 0.264 (NS) |
| low performer_Peak | 0.702 (NS) | 0.572 (NS) | 0.508 (NS) | 0.249 (NS) |
| middle performer_LME | 0.160 (NS) | 0.637 (NS) | 0.102 (NS) | 0.516 (NS) |
| middle performer_Peak | 0.132 (NS) | 0.833 (NS) | 0.059 (NS) | 0.237 (NS) |
| high performer_LME | 0.103 (NS) | 0.740 (NS) | 0.098 (NS) | 0.360 (NS) |
| high performer_Peak | 0.049 (*) | 0.460 (NS) | 0.088 (NS) | 0.325 (NS) |

### Time-Point–Specific Divergence

#### Clock^*Out*^ mutants exhibited impaired sustained climbing, with deficits persisting and intensifying over time in males, while females showed milder or age-stable effects compared to *w*^1118^

Presenting the heatmaps of –log_10_(p) transformation instead of just the raw p-values shows enhanced perceptual spacing since raw p-values tend to cluster tightly between 0 and 0.05 which could be compressing meaningful differences into a narrow band of values. Transforming with –log_10_spreads them non-linearly making gradations of significance much easier to visually distinguish. This handles a well-known limitation in heatmap color perception. Color scales that are tied to −log_10_(p) emphasize high-significance time points as intense “hot spots” thus enhancing visual salience of significant results. In the resulting color scale, low significance points (p ≈ 0.1– 0.05) appear as faint warm tones (–log_10_≈ 1–1.3), moderate significance (p ≈ 0.05–0.01) as brighter yellows (– log_10_≈ 1.3–2), and high significance (p ≤ 0.001) as intense reds (–log_10_≥ 3). For males, wk1 Clock^*Out*^ mutants initiate climbing similarly to the wk1 *w*^1118^ controls but rapidly lose pace after 8 seconds. This indicates that clock disruption impairs sustained upward locomotion in the mid-phase of the assay. The wk3 Clock^*Out*^ mutants exhibit a slightly delayed onset of impairment compared to wk1 genotype comparison. This indicates that the effect of clock disruption on geotaxis persists and intensifies. For aging effects, *w*^1118^ controls showed remarkably stable climbing kinetics between wk1 and wk3, indicating minimal aging-related decline in this time window. The Clock^*Out*^ mutants show a modest additional slowing with age potentially hinting at cumulative impairment. The effects are stronger than *w*^1118^ aging but are weaker than in genotype comparisons. For females, the wk1 Clock^*Out*^ mutants show a delayed but detectable deficit in climbing height compared to wk1 *w*^1118^ controls. The wk3 Clock^*Out*^ mutants remain statistically indistinguishable from wk3 *w*^1118^ peers in time-resolved peak height. For aging effects, the *w*^1118^ controls display reduced initial climbing performance with age, climbing more slowly in the first few seconds at wk3 versus wk1, but then converge in the sustained phase. The Clock^*Out*^ mutants remained phenotypically consistent between wk1 and wk3, suggesting that their plateaued performance at wk1 carries forward, and aging contributes little beyond the initial clock-loss deficit.

#### Overall Summary and Insights

For genotype effects on males at both wk1 and wk3, Clock^*Out*^ males climb significantly lower than *w*^*1118*^ controls and produce markedly fewer high performers. Their deficit appears in the mid-phase of the assay (8–10s), where sustained climbing breaks down for the Clock^*Out*^ males. Clock^*Out*^ females showed a small but significant drop in peak height only at wk1. By wk3, they are indistinguishable from w^1118^ controls at all time points and in all performer categories. For aging effects on males, there was no statistical change in w^1118^ controls for overall climbing trajectory, peak height, or any performer categories suggesting that climbing remained stable between wk1 and wk3. For Clock^Out^ males, there was a small decline in climbing trajectory from wk1 to wk3, though less pronounced than the genotype effect itself. For female aging effects on w^1118^ controls, there was a modest early-phase decline in climb and fewer high performers at wk3 flies, but convergence with wk1 controls in later time bins. For Clock^Out^ mutant females, there was no change with age as performance at wk3 mirrors the plateau established at wk1. Instantaneous climbing velocity is unaffected by genotype or age in both sexes indicating that clock disruption and mild aging impair sustained height gain without altering instantaneous climbing velocity.

The automated geotaxis pipeline presented in this paper was able to extract that for males, *w*^1118^ controls had stable geotaxis performance across age with high consistency in climbing height and high performer rates and Clock^*Out*^ mutants had clear and consistent climbing impairment at both ages. Aging compounds these deficits, especially in trajectory measures and time-specific divergence. These flies climb normally at first but cannot sustain upward motion after ∼8–9 seconds. For females, *w*^1118^ controls showed aging reduced early climbing effort (0–4 s), but performance stabilizes mid-assay. High performers remain consistent. The Clock^*Out*^ mutants had mild genotype effect at wk1 (reduced peak height), but performance converges with *w*^1118^ by wk3. Aging has a minimal additional impact. The clock gene disruption impairs sustained geotaxis, especially in males, and these deficits worsen with age. Females are less sensitive overall, and aging appears to selectively affect early-phase performance rather than total climbing ability.

## 4. Discussion

### Significance of the Developed Device

The geotaxis assay platform we have engineered represents a leap forward in both precision and scalability for *Drosophila* locomotor studies. Physically, our 3D-printed vial rack and third-class lever tapping mechanism (**Supplementary Figure 1b**) enforce identical starting conditions across all 12 vials. Illumination is standardized via a fixed light source, which eliminates glare and shadows that can confound background subtraction. Computationally, our end-to-end pipeline, from video conversion to vial detection to statistical modeling (**Figure 2b**), removes all human intervention once metadata files are created for each video. By embedding LME model fitting and non-parametric tests directly into the workflow, days of manual scoring and offline analysis turn into just hours of unsupervised processing. The device’s modular architecture and PRISM-friendly output CSVs further ensure that labs can swap in custom camera modules or repeat the assay under altered environmental settings (temperature, humidity) without rewriting code. Taken together, this system not only removes observer bias and processing time by nearly threefold (**Supplementary Figure 2a–b**) but also opens access to high-resolution, reproducible locomotor phenotyping across research groups of various computational capabilities.

### Impact of Advanced Behavioral Analysis

This pipeline generates on the order of 10^5^ centroid position measurements per video folder which is roughly 779 × more data than a single manual frame count (**Supplementary Figure 2b**). This explosion in data density enables two transformative advances. First, zoning each fly as a low, middle, or high performer in real time (**Figure 3c**) captures within-population heterogeneity that masked subtler phenotypes in prior assays. Second, continuous time-series analyses of position, velocity, and zoning metrics feed directly into mixed-effects models. This yields in both genotypes (or aging) main effects and precise genotypes (or aging) × time interactions (**Table 1**). The resulting heatmaps of –log_10_(p) (**Figure 5c, f, i, l**) pinpoint the exact seconds when a comparison lineage diverges. While GPU-powered vial detection accelerates processing, we are already prototyping rule-based contour detection and pruned, mobile-optimized neural networks to bring this capacity to laptops and Raspberry Pis themselves, ensuring that small labs and teaching environments can adopt the full pipeline without access to large computer resources.

### Biological Implications of Machine Learning Platform

Compared with age matched control, or during age, *Clock*^*Out*^ male showed a severe climbing deficit (**Figure 5 a,d**; **Table 1a**) reflects impaired motor coordination rather than energy depletion. Mid-assay breakdown (8–10 s) occurs without velocity changes, indicating intact movement initiation but compromised endurance. However, in female phenotypes, *Clock*^*Out*^ females show mild, age-resolving deficits: wk1 peak height reduction normalizes by wk3 (Table 1b). Also, when compared with age matched control *Clock*^*Out*^ females mild phenotypes, suggesting that sex specific difference possible due to different metabolic rate in *Clock*^*Out*^.

Glaz driven knockout of *PolG* males exhibit age-amplified decline (**Supplementary Figure 4, Supplementary Table 1a**), with V106955 alleles showing significant wk7 trajectory loss and increased low performers. Elav PolG males display progressive deterioration (**Supplementary Figure 5, Supplementary Table 2a**), particularly in V106955 at wk7, confirming mitochondrial dysfunction exacerbates aging. Glaz PolG females demonstrate resilience but not net improvement: aged (wk7) mutants outperform controls in climbing height (**Supplementary Table 1b**) but exhibit significant age-related decline. Elav PolG females exhibit delayed dysfunction, not enhancement: transient wk3 preservation (**Supplementary Table 2b**) precedes wk7 decline, indicating postponed mitochondrial pathology rather than true beneficial effects.

**Table 2.**
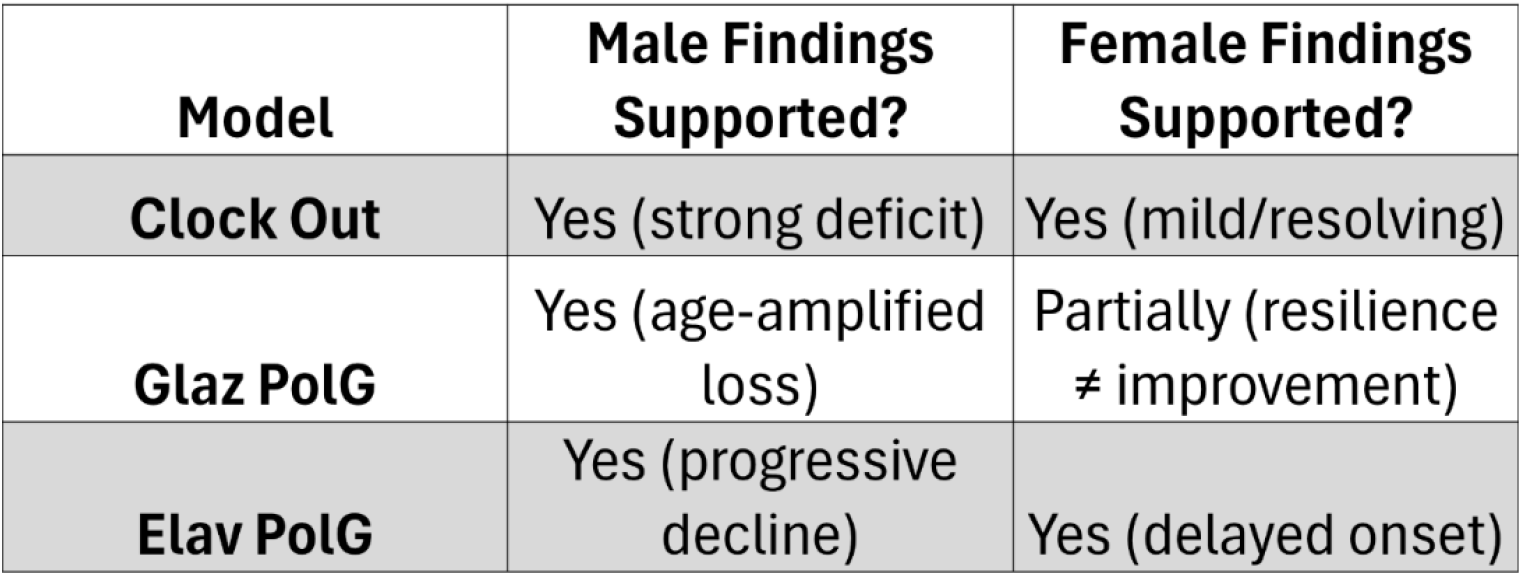
Sex-Specific Validity of Biological Insights: Whether Clock^Out^, Glaz>UAS-PolG^RNAi^, and Elav>UAS-PolG^RNAi^ locomotor phenotypes are fully supported in males and females. “Yes” denotes validated significant effects; “Partially” indicates qualified resilience rather than true improvement.

The sex-dimorphic trajectories (**Supplementary Tables 1–2**) reveal fundamental neuromuscular aging principles. Male vulnerability in clock/mitochondrial mutants parallels human Parkinson’s and POLG-ataxias, where men show earlier onset. Female resilience, notably in Glaz PolG, aligns with estrogen’s neuroprotection via PGC-1α-mediated mitochondrial biogenesis. The dissociation between intact velocity (brainstem/spinal circuits) and impaired climbing persistence (vulnerable dopaminergic pathways) underscores hierarchical motor control (Figure 4). Transient benefits in young PolG females resemble “mitohormesis,” where mild oxidative stress upregulates cytoprotective pathways, highlighting therapeutic potential for sex-stratified neurodegenerative treatments. This platform bridges Drosophila genetics to conserve sexual dimorphism in aging, emphasizing the necessity of studying both sexes in translational research.

### Future Directions

Looking ahead, we envision several key enhancements to broaden the assay’s capabilities and accessibility. First, by deploying compact, on-device inference engines, such as TensorRT-pruned Faster R-CNN models or efficient rule-based heuristics, directly on Raspberry Pi 4B units, we can enable live vial detection, tracking, and preliminary statistical summaries without reliance on external GPUs. Second, integrating additional camera angles and depth sensors (stereo or structured light) will allow us to capture flight kinematics, courtship rituals (wing extension, chasing), and full 3D locomotion, thereby enriching phenotype catalogs for social and sensorimotor research. Third, adding programmable light, temperature, and humidity controllers alongside LED-driven optogenetic stimulation will permit tightly synchronized environmental and neural activation paradigms, offering new insights into behavior under dynamically controlled conditions. Fourth, adapting the vial holder to a 96-well plate format, with automated fly loading via micro-aspirators and robotic liquid handlers for drug or RNAi administration, will transform the platform into a high-throughput screening workhorse for chemical libraries or genome-wide knockdown collections. Finally, we plan to launch a cloud-based repository and web portal hosting pre-trained models, standardized datasets, and interactive dashboards, complete with plug-and-play modules for novel behaviors and statistical analyses, thereby fostering a collaborative community ecosystem for continued innovation and comparative studies.

## 5. Conclusion

Our integrated hardware-software geotaxis assay unifies standardized device engineering, deep-learning vial segmentation, and high-resolution computer-vision tracking into a turnkey platform that outstrips manual methods in speed (2.8× faster), data volume (≈779× richer), and analytical sophistication. By resolving continuous, zone-specific locomotor metrics and embedding robust statistical modeling, we explain subtle, sex- and age-dependent phenotypes across circadian and mitochondrial mutant lines, with these insights previously obscured by coarse endpoint scoring. The device’s modularity, user-friendly GUI (**Figure 5**), and open-format outputs position it for rapid adaptation to new behavioral paradigms, from courtship to optogenetic stimulation. Looking forward, implementing real-time on-device analytics, environmental control, and ultra-high-throughput screening designs would extend its reach, while a community platform for sharing models and data will accelerate discovery across Drosophila and broader neurobehavioral research. In sum, this work establishes a new standard for automated behavioral phenotyping, bridging technical accessibility with powerful, nuanced biological insight.

## Acknowledgments

We are grateful for access to the University of Alabama at Birmingham’s High-Performance Computing (Cheaha) resources, ensuring efficient processing of large-scale video datasets.

## Supporting information Text

Supplementary Figure 1. Design and Assembly of the Automated Geotaxis Assay Device

Supplementary Figure 2. Device Performance Metrics vs. Manual Assay

Supplementary Figure 3. IoU Distribution for Deep Learning Performance

Supplementary Figure 4. Glaz-driven PolG RNAi lines Geotaxis Behavior and Statistical Analysis Visualizations

Supplementary Figure 5. Elaz-driven PolG RNAi lines Geotaxis Behavior and Statistical Analysis Visualizations

Supplementary Tables 1. P-values for trajectory and peak metrics comparing Glaz-driven PolG RNAi lines

Supplementary Table 2. P-values for trajectory and peak metrics comparing Elav-driven PolG RNAi lines

## Author contributions

DM outlined the manuscript under GCM supervision and conducted all the machine learning experiments, generated the figures and wrote the paper. NM made the app and generated the figure. DP did all the genetic experiments and SD prepared the locomotor device using 3D printing. Conduction. All the authors reviewed the manuscripts. GCM is also involved with conceptualization, data curation, funding acquisition: writing/editing and project administration.

## Data and Code availability

The copyright codebase as well as example notebooks for analysis are being prepared and will be made publicly available on GitHub under a GPL2.0 license upon acceptance.

## References

1. Ali, Y. O., Escala, W., Ruan, K., & Zhai, R. G. (2011). Assaying locomotor, learning, and memory deficits in Drosophila models of neurodegeneration. Journal of Visualized Experiments, 49. 10.3791/2504

2. Aggarwal, A., Reichert, H., & VijayRaghavan, K. (2019). A locomotor assay reveals deficits in heterozygous Parkinson’s disease model and proprioceptive mutants in adult Drosophila. Proceedings of the National Academy of Sciences of the United States of America, 116(49). 10.1073/pnas.1807456116

3. Branson, K., Robie, A. A., Bender, J., Perona, P., & Dickinson, M. H. (2009). High-throughput ethomics in large groups of Drosophila. Nature Methods, 6(6). 10.1038/nmeth.1328

4. Gargano, J. W., Martin, I., Bhandari, P., & Grotewiel, M. S. (2005). Rapid iterative negative geotaxis (RING): a new method for assessing age-related locomotor decline in Drosophila. Experimental Gerontology, 40(5), 386–395. 10.1016/J.EXGER.2005.02.005

5. Cao, W., Song, L., Cheng, J., Yi, N., Cai, L., Huang, F. de, & Ho, M. (2017). An Automated Rapid Iterative Negative Geotaxis Assay for Analyzing Adult Climbing Behavior in a Drosophila Model of Neurodegeneration. Journal of Visualized Experiments : JoVE, 2017(127), 56507. 10.3791/56507

6. Liu, H., Han, M., Li, Q., Zhang, X., Wang, W. A., & Huang, F. de. (2015). Automated rapid iterative negative geotaxis assay and its use in a genetic screen for modifiers of Aβ42-induced locomotor decline in Drosophila. Neuroscience Bulletin, 31(5), 541. 10.1007/S12264-014-1526-0

7. Spierer, A. N., Yoon, D., Zhu, C. T., & Rand, D. M. (2021). FreeClimber: Automated quantification of climbing performance in Drosophila. Journal of Experimental Biology, 224(2). 10.1242/JEB.229377/267839/AM/FREECLIMBER-AUTOMATED-QUANTIFICATION-OF-CLIMBING

8. Canic, T., Lopez, J., Ortiz-Vega, N., Zhai, R. G., & Syed, S. (2025). High-resolution, high-throughput analysis of Drosophila geotactic behavior. Journal of Experimental Biology, 228(4). 10.1242/JEB.248029/VIDEO-2

9. Livelo, C., Guo, Y., Madhanagopal, J., Morrow, C., & Melkani, G. C. (2025). Time-restricted feeding mediated modulation of microbiota leads to changes in muscle physiology in Drosophila obesity models. Aging Cell, 24(2). 10.1111/acel.14382

10. Pasam, E. S., Madamanchi, K., & Melkani, G. C. (2025). Dissecting metabolic regulation of behaviors and physiology during aging in Drosophila. Biogerontology, 26(5). 10.1007/s10522-025-10306-y

11. Edwards, L. J., Muller, K. E., Wolfinger, R. D., Qaqish, B. F., & Schabenberger, O. (2008). An R2 statistic for fixed effects in the linear mixed model. Statistics in Medicine, 27(29). 10.1002/sim.3429

12. Gibbons, R. D., Hedeker, D., & Dutoit, S. (2010). Advances in analysis of longitudinal data. In Annual Review of Clinical Psychology (Vol. 6). 10.1146/annurev.clinpsy.032408.153550

13. Verbeke, G., Molenberghs, G., & Rizopoulos, D. (2010). Random effects models for longitudinal data. In Longitudinal Research with Latent Variables. 10.1007/978-3-642-11760-2_2

14. West, B. T., Welch, K., Galecki, A., & Gillespie, B. (2022). Linear mixed models: A practical guide using statistical software. In Linear Mixed Models: A Practical Guide Using Statistical Software. 10.1201/9781003181064

15. Wilson, D. J. (2019). The harmonic mean p-value for combining dependent tests. Proceedings of the National Academy of Sciences of the United States of America, 116(4). 10.1073/pnas.1814092116

